# Genome sequence and RNA-seq analysis reveal genetic basis of flower coloration in the giant water lily *Victoria cruziana*

**DOI:** 10.1101/2024.06.15.599162

**Authors:** Melina Sophie Nowak, Benjamin Harder, Samuel Nestor Meckoni, Ronja Friedhoff, Katharina Wolff, Boas Pucker

## Abstract

*Victoria cruziana* is well known for its huge floating leaves covered with sharp spines and its night blooming. Reports indicate that white flowers open during the first night and turn light pinkish during the following day and the second night. Here, we set out to unravel the molecular basis and ecological function of the flower color change in *V. cruziana*. A high quality genome sequence with a contig N50 of 44.2 Mbp, a scaffold N50 of 300 Mbp, and a total assembly size of 3.5 Gbp was generated as the genetic basis for this study. Comparative transcriptomics revealed the genes required for anthocyanin biosynthesis genes and their transcriptional regulators as differentially expressed between the white and the light pinkish stage of a flower. Structural genes with expression differences between white and light pinkish flower stages include *VcrF3’H*, *VcrF3’5’H*, *VcrDFR*, *VcrANS*, and *VcrarGST*. The expression pattern of the corresponding transcription factors *VcrMYB123*, *VcrMYB-SG6_a*, *VcrMYB-SG6_b, VcrTT8*, and *VcrTTG1* also showed differences that aligned with the flower color.

## Introduction

Waterlilies of the genus *Victoria* are best known for their giant floating leaves that can endure a substantial weight if equally distributed across the leaf. The genus *Victoria* belongs to the *Nymphaeaceae* and harbors three robust species *V. amazonica*, *V. cruziana* and *V. boliviana* that were recently proposed (Smith *et al*., 2022). These aquatic species are night bloomers, considered as short-lived perennial plants and naturally grow under tropical conditions. Distinctive shared characteristics are the sharp spines covering most plant parts, massive leaves, and a grand flower. The whole development from seeds to a mature giant water lily takes less than five months (Smith *et al*., 2022). *Victoria spp.* also have an economic relevance, as the large seeds and rhizomes can be used as a food source (Bortolotto *et al*., 2015).

The flower, a special feature of *Victoria spp.,* has been reported to undergo a color change from white to light pinkish (**Figure 1**) (Wu *et al*., 2018). Development of the *Victoria* flower bud starts underwater and the bud only opens after reaching the water surface. Interestingly, *Victoria* plants bloom at night. To the best of our knowledge, one individual plant exhibits only a single blooming flower at any given time. Each flower undergoes a flowering period of two consecutive nights, during which the flower color changes from white to light pinkish. In addition, anthesis also occurs during these consecutive nights, starting with the female reception, followed by staminate maturation (male phase) (Seymour & Matthews, 2006; Jiang *et al*., 2021). The ecological reason and molecular basis of the flower color change are currently unknown, but multiple hypotheses exist that could explain it: (1) flower color might change following a visitation/pollination event, (2) light could trigger the accumulation of pigments once flowers emerge from the water, or (3) a developmental program could lead to pigmentation in an age-dependent manner. In the following sections, we will provide context for each of these hypotheses.

**Figure 1:**
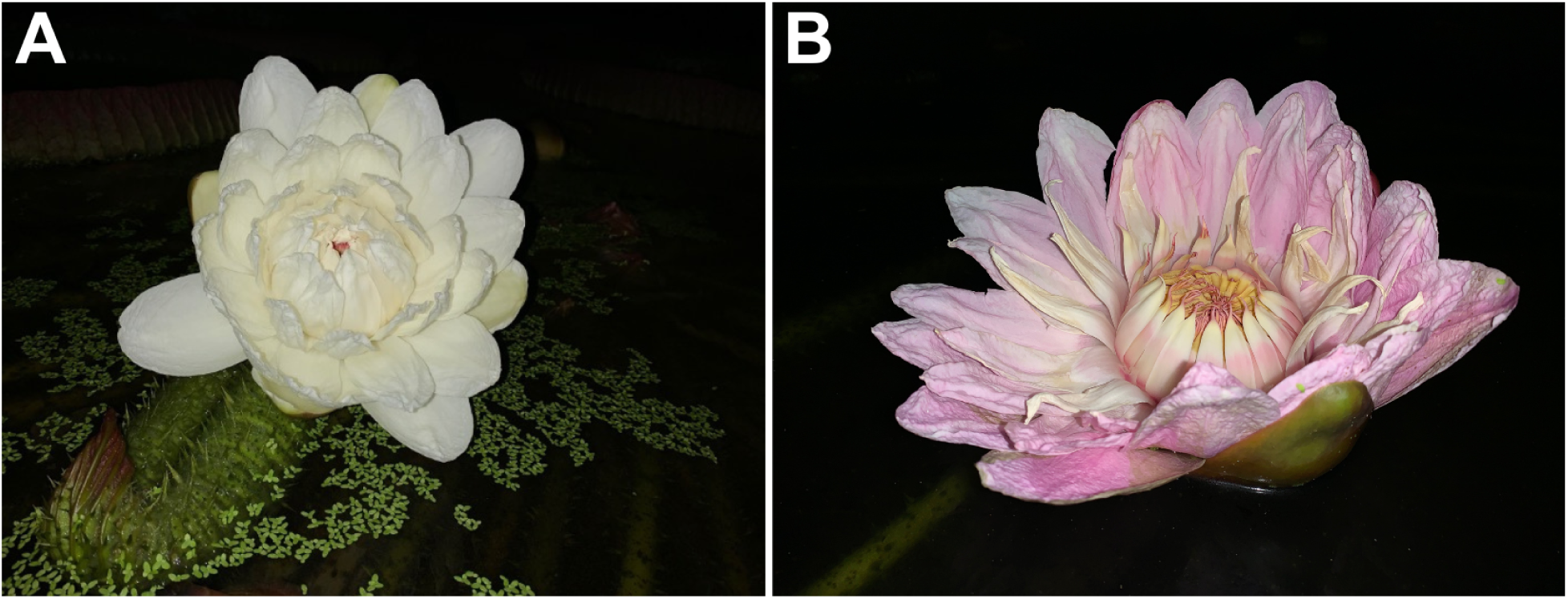
Flower of *Victoria cruziana*. Comparison of a flower between the first bloom (white) at night (A) and the consecutive night (light pink) (B).

Previous studies described flower color changes following pollination events for other plant species (Weiss, 1991; Ruxton & Schaefer, 2016). Although these events are rare, floral color change can serve as a signal between plant and insect to transmit the information that pollination, or at least visitation, already occurred (Weiss, 1991; Ruxton & Schaefer, 2016; Garcia *et al*., 2022). Flowers of *V. cruziana* appear white in the first night, combined with a fruity smell, secreted by the carpellary appendages which could suggest a function in pollinator attraction (Zini *et al*., 2019). A previously reported pollinator of *Victoria spp.* are *Cyclocephalini,* a tribe of scarab beetles that are pantropical distributed (Moore & Jameson, 2013). Willmer et al. described the floral color alterations from lilac to white in *Desmodium setigerum* after receiving visitors (Willmer *et al*., 2009).

Light-induced accumulation of red anthocyanins has been described before in many plant species like *Arabidopsis thaliana* (Maier *et al*., 2013; Li *et al*., 2016), *Malus spec.* (Merzlyak & Chivkunova, 2000; Takos *et al*., 2006), or *Pyrus spec.* (Dussi *et al*., 1995). The molecular basis of this light response has been extensively studied and involves an up-regulation of the transcription factors (TFs) controlling the anthocyanin biosynthesis genes (Takos *et al*., 2006; Li *et al*., 2016). Anthocyanins can serve as sunscreen thus protecting plants against excessive light irradiation (Gould, 2004) and are strong antioxidants that can scavenge reactive oxygen species (Wang *et al*., 1997).

Anthocyanin formation might also take place as part of a developmental program, thus depending on the age of the flower and not directly connected to environmental conditions. For example, a study in *Fuchsia excorticata* demonstrated that the flower color change from green to red is age-dependent and not caused by pollination (Delph & Lively, 1989). An investigation in *Pulmonaria collina* also noted an age-dependent color change from red to blue and explains this as a mechanism to direct pollinators towards the young flowers thus increasing the pollination efficiency (Oberrath & Böhning-Gaese, 1999). Similar findings have been reported for *Pedicularis monbeigiana* which shows a color change from white to purple with increasing flower age (Sun *et al*., 2005).

The pigments responsible for flower coloration are well studied and often belong to the anthocyanins, carotenoids, or betalains (Tanaka *et al*., 2008). Anthocyanins and carotenoids are taxonomically widespread and responsible for most flower colors (Tanaka *et al*., 2008), while betalains are restricted to the Caryophyllales (Timoneda *et al*., 2019). Anthocyanins are derived from phenylalanine involving the general phenylpropanoid pathway and the flavonoid biosynthesis (**Figure 2**). Structural genes required for the anthocyanin biosynthesis include *phenylalanine ammonia lyase* (*PAL*), *cinnamate 4-hydroxylase* (*C4H*), and *4-coumarate-CoA ligase* (*4CL*) as part of the general phenylpropanoid pathway followed by *chalcone synthase* (*CHS*), *chalcone isomerase* (*CHI*), *flavanone 3-hydroxylase* (*F3H*), *dihydroflavonol 4-reductase* (*DFR*), *anthocyanidin synthase* (*ANS*), *anthocyanin-related Glutathione S-transferase* (*arGST*), and *UDP-glucose: flavonoid-3-O-glucosyltransferase* (*UFGT*) as one branch of the flavonoid biosynthesis (Winkel-Shirley, 2001; Eichenberger *et al*., 2023). Activity of these structural genes is orchestrated by an ensemble of transcription factors. The two largest transcription factor families in plants, MYBs and bHLHs, contribute substantially to the regulation of different branches of the flavonoid biosynthesis. MYB transcription factors are able to bind DNA through a highly conserved DNA-binding repeat domain (Prouse & Campbell, 2012), while bHLHs can bind DNA with a stretch of basic amino acids (Voronova & Baltimore, 1990). MYBs can be divided into different groups based on the number of characteristic repeats with the R2R3-MYBs playing a predominant role in the control of the flavonoid biosynthesis (Baudry *et al*., 2004; Stracke *et al*., 2007; Gonzalez *et al*., 2008; Li, 2014; Marin-Recinos & Pucker, 2024). Specific subgroups of these R2R3-MYBs have different and even opposing functions. Subgroup 4, including *AtMYB4*, *AtMYB32*, *AtMYB7*, *AtMYB3*, is known for encoding transcription factors with repressive functions (Jin *et al*., 2000; LaFountain & Yuan, 2021). *AtMYB123* and *AtMYB5* belong to subgroup 5 and regulate proanthocyanidin biosynthesis in *Arabidopsis thaliana* (Baudry *et al*., 2004; Xu *et al*., 2014). MYBs assigned to subgroup 6 (*AtMYB113, AtMYB114, AtMYB90, AtMYB75*) are involved in controlling the anthocyanin biosynthesis in *A. thaliana* (Borevitz *et al*., 2000; Marin-Recinos & Pucker, 2024) whereas *AtMYB111, AtMYB12 and AtMYB11* (subgroup 7) are regulating the production of flavonols (Stracke *et al*., 2007; Dubos *et al*., 2010). Over the last years, various orthologous of the well characterized MYBs in *A. thaliana* have been identified across plant species which enables a broader understanding of regulatory mechanisms in the plant kingdom. For example, *At*MYB123 orthologs have been reported to control the anthocyanin biosynthesis in addition to the proanthocyanidin biosynthesis (Martínez-Rivas *et al*., 2023) and orthologs of the *At*MYB5 have been reported to activate the anthocyanin biosynthesis in strawberry (Jiang *et al*., 2023). This suggests that the generally well conserved regulation of the anthocyanin biosynthesis might be subject to lineage-specific differences. Given that *A. thaliana* does not have colorful fruits and flowers, it is likely that many secrets of the anthocyanin regulation can only be discovered by exploring non-model species.

**Figure 2:**
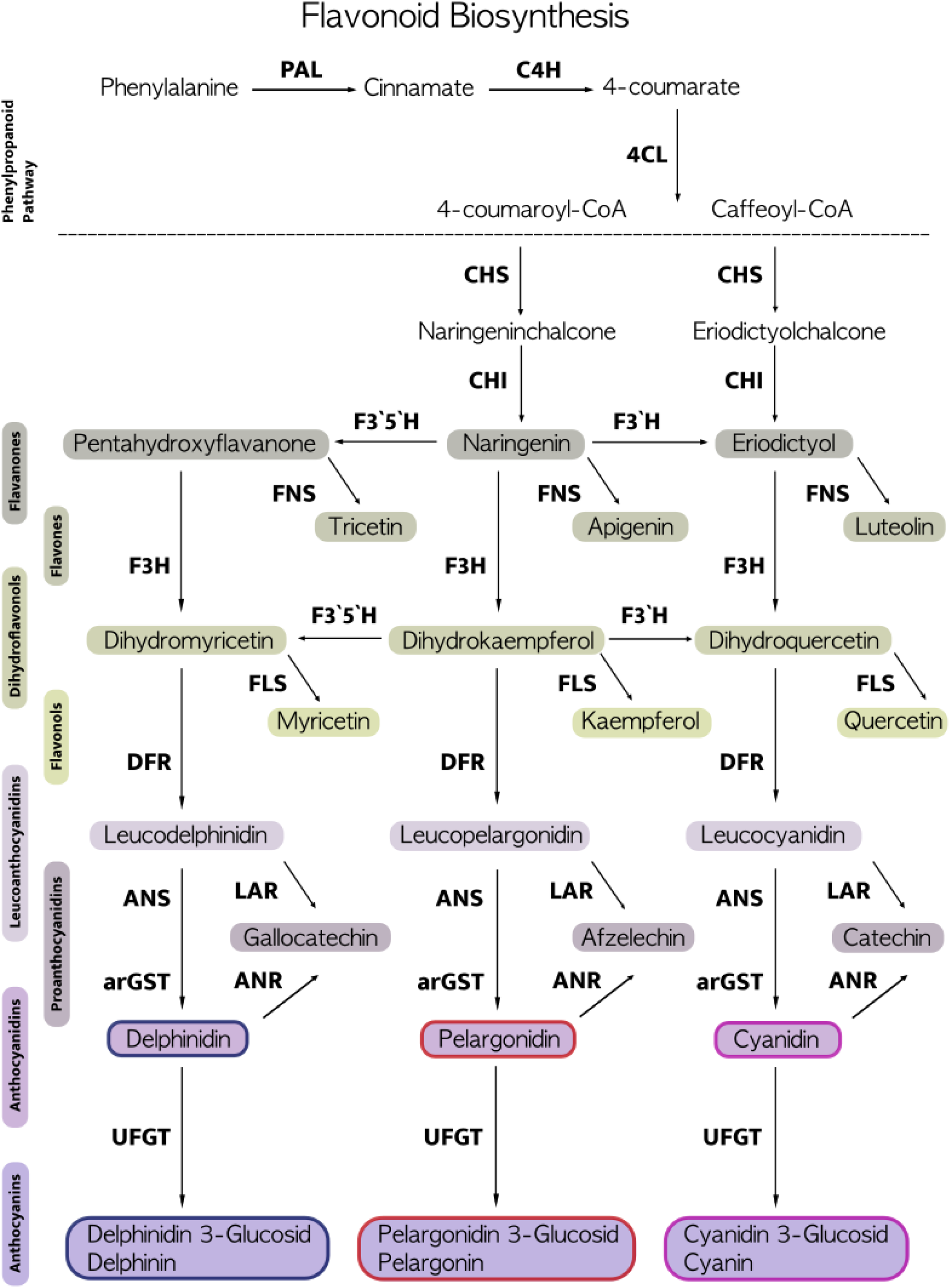
Schematic illustration of the phenylpropanoid and flavonoid biosynthesis pathway including the reactions leading to anthocyanins. PAL (Phenylalanine ammonia lyase), C4H (Cinnamate 4-hydroxylase), 4CL (4-coumarate-CoA ligase), CHS (Chalcone synthase), CHI (Chalcone isomerase), FNS (Flavone synthase), F3H (Flavanone 3-hydroxylase), F3’H (Flavonoid 3’-hydroxylase), F3’5’H (Flavonoid 3’,5’-hydroxylase), FLS (Flavonol synthase), DFR (Dihydroflavonol 4-reductase), LAR (Leucoanthocyanidin reductase), ANR (Anthocyanidin reductase), ANS (Anthocyanidin synthase), arGST (Anthocyanin-related Glutathione S-transferase), UFGT (UDP-glucose: flavonoid-3-O-glucosyltransferase).

Previous studies investigated the pigment composition of *Victoria cruziana* and identified four anthocyanins (two cyanidin and two delphinidin derivatives) in the flower (Wu *et al*., 2018). An increase in anthocyanin content across the blooming days correlates with the intensifying pigmentation and supports the role of anthocyanins as the predominant flower colorants (Wu *et al*., 2018).

To identify the genes involved in the anthocyanin biosynthesis and its regulation in *V. cruziana*, we performed long-read sequencing of the genome and transcriptome. Here, we provide a genome sequence of *V. cruziana* and a corresponding annotation supported by RNA-seq and direct RNA sequencing that serve as the basis for metabolic pathway explorations. Additionally, we conducted RNA-seq experiments to identify differentially expressed genes between white and light pinkish flowers in two developmental stages of *V. cruziana*. Specifically, we aimed to unravel the genetic network underlying the flower color transition from white to light pink and to set our findings in an ecological context.

## Material and Methods

The genome of *V. cruziana* was sequenced with Oxford Nanopore Technologies (ONT) long reads to generate a high quality genome sequence that served as basis for the transcriptomics investigation based on direct RNA sequencing and RNA-seq (**Figure 3**).

**Figure 3:**
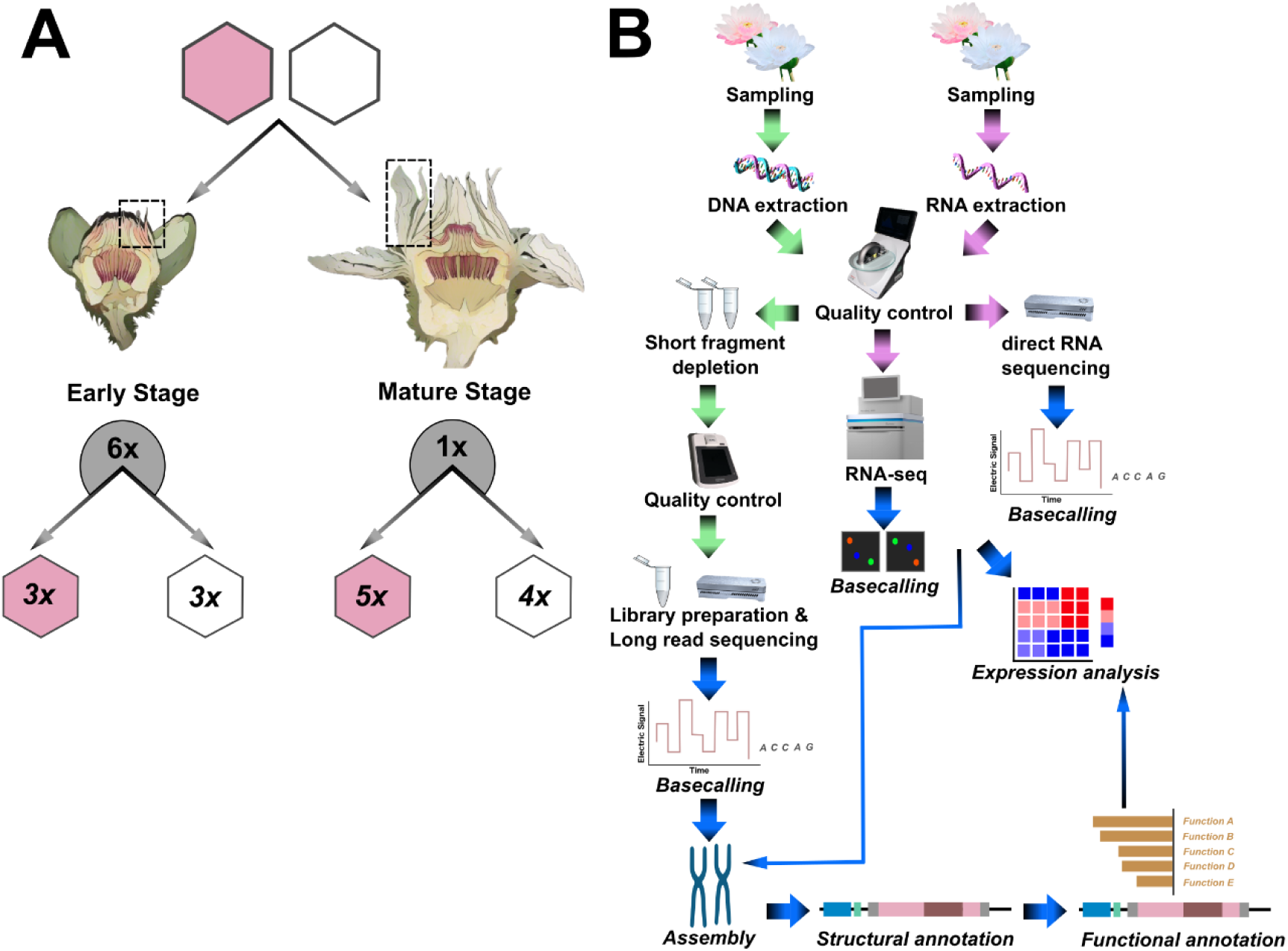
Experimental design (A) and workflow for genomic and transcriptomic analysis (B) of flower color in *Victoria cruziana.* Flower color of white and light pink petals has been comparatively analyzed in (I) an early stage of *V. cruziana* ranging from buds between day 3 and day 8 and in (II) the mature stage as the flower fully opened. In total 6 individual flower buds of the early stage have been sampled whereas 1 individual flower bud was used for analysis in the mature stage (A). Green arrows within the workflow depicts DNA related wet lab procedures whereas pink represents RNA related wet lab. Blue arrows indicate all bioinformatic related analyses.

### Plant cultivation, DNA extraction and RNA extraction

For genomic DNA extraction, a *V. cruziana* plant (XX-0-BRAUN-7477852) was cultivated at 28 °C in a 500 L container to facilitate repeated sampling of young leaves and immediate processing of material. Light was provided for 16 hours per day with two LEDs (Niello^®^, 300W) placed 1 m above the water surface. High molecular weight DNA extraction and quality assessment were performed based on a previously developed CTAB protocol optimized for high molecular weight DNA extraction from plants (Siadjeu *et al*., 2020; Wolff *et al*., 2023). In brief, fresh leaf material was ground in liquid nitrogen and the fine powder subjected to the CTAB-based DNA extraction procedure. RNase treatment in the CTAB-TE buffer was performed overnight at room temperature. After initial quality assessment via NanoDrop and agarose gel, short DNA fragments were depleted with the Short Read Eliminator kit. DNA quantification prior to library preparation for the nanopore sequencing was performed via Qubit.

For transcriptome analysis, young and mature *V. cruziana* petals at the white and light pink coloring stage were collected. Sampling of mature samples has been conducted in two consecutive nights during July 2023 in a dedicated *Victoria* glasshouse located in the Botanical Garden of TU Braunschweig (52.27043343004546, 10.5336707275785). Samples of young petals have been collected during September and October of 2024. Petals of the same flower at both color and developmental stages were carefully detached, immediately frozen in liquid nitrogen, and stored at -70 °C prior to RNA extraction. The initial sampling for mature petals occurred on July 19th, 2023, at 10:50 pm. Four samples were taken from the not yet fully opened white flower. The weather on this day was cloudy. The subsequent day, July 20th 2023, had cloudy and sunny conditions, and five samples were collected from the light pink blooming flower. The ambient temperature in the greenhouse was 24 °C during the day and 22 °C at night. Sampling of young flower buds has been performed on September 20th, September 24th, and October 4th at around 6:30 pm. The ventilation temperature in the greenhouse remained constant at 34 °C, while humidity fluctuated between 80 % and 100 %. The plants did not receive any artificial light exposure. For irrigation, tap water with a temperature of 27 °C was used. The plants were cultivated in a pool with a depth of 1.07 meters and a width of 8.8 m x 5.5 m, with a substrate ratio of 1:1 (compost to peat). Each liter of water received 6 grams of Osmocote Pro and 6 grams of lime as fertilizer.

Total RNA of young and mature petals was extracted with the RNA Plant and Fungi kit (Macherey & Nagel), together with a DNase treatment according to the manufacturer’s instructions. Four young petals buds between day 3 and day 8 have been harvested and petals of the opened flower from the mature stage. Cell homogenization was performed using the Bertin Ribolyser at 6000 RPM with three cycles of 30 seconds each, with a 30 second break between cycles, using 3 mm steel beads. The extracted RNA quality and quantity was determined by a NanoDrop measurement and an agarose gel. The RNA was stored at -70 °C until it was sent for RNA-seq. Paired-end RNA-seq (2 x 150 nt) was conducted using Illumina NovaSeq 6000 (BMKGene).

### Pore-C

For Pore-C, fresh leave samples from a *V. cruziana* plant (XX-0-BRAUN-7477852) cultivated at 28 °C in a 500 L container with light periods for 16 hours per day has been utilized. The Pore-C workflow was performed as described in the published adapted protocol for *Victoria cruziana* (Nowak & Pucker, 2025), that is based on several previously published protocols (Deshpande *et al*., 2022). Briefly, the protocol is divided across three days for the main laboratory part, followed by size selection, library preparation, and sequencing. Day one comprised crosslinking, cryogrinding, nuclei isolation and chromatin denaturation. Crosslinking was achieved through a vacuum dependent infiltration of formaldehyde with glycine administration to terminate the crosslinking reaction. Cryogrinding has been performed with liquid nitrogen followed by subsequent nuclei permeabilization and isolation. For chromatin denaturation the restriction enzyme Nlall was used, together with a heat-based inactivation on day two. Proximity ligation was conducted using T4 DNA ligase where the cohesive ends have been ligated into chimeric polymers. After the ligation step was finished, protein degradation with proteinase K and SDS was performed. During an overnight incubation at 56°C de-crosslinking took place. On day three, a phenol-chloroform based DNA extraction was carried out with subsequent ethanol precipitation. To enhance DNA precipitation, 5 M NaCl and 3 M sodium acetate pH 5.5 were added to the obtained chimeric dsDNA mixture. Extracted DNA has been measured with a qubit utilizing the dsDNA BR Assay. Afterwards the sample was stored at 4°C prior to size selection and library preparation.

To specifically enrich obtained Pore-C fragments greater than 2 kb, a modified size selection has been performed. This method is based on solid-phase reversible immobilization (SPRI) which was originally published by Schalamun & Schwessinger (Schalamun & Schwessinger, 2017). The herein utilized custom size selection buffer contained AMPure XP beads and a specified concentration of 40% (w/v) PEG 8000 to determine the recovered fragment length. For maximized chimeric Pore-C fragment output, DNA was diluted to 60 ng/µl in 50 µl TE buffer pH 8.0 and 0.85X volume of the custom prepared buffer was used to perform size selection (Nowak & Pucker, 2025). Size selected DNA was again quantified using a Quibit and stored at 4°C until library preparation.

### Genome sequencing

Sequencing was performed on R9 and R10 flow cells. Per library for R9 flow cells, 1 µg of high molecular weight DNA was utilized following ONT’s SQK-LSK109 protocol. Priming of R9.4.1 flow cells and washing steps between sequencing runs were performed according to ONT’s protocols (EXP-FLP002, EXP-WSH004). Sequencing was performed on a MinION Mk1B. FAST5 files were collected and subjected to basecalling using Guppy v6.4.6+ae70e8f with the dna_r9.4.1_450bps_hac model and a minimum quality score of 9. Library construction for R10 flow cells was performed using 1µg of high molecular weight DNA following the SQK-LSK114 protocol. Priming and washing of the flow cells was performed as described above. Pore-C library preparation was identically performed as for long-read sequencing besides a minor variation in the DNA repair and end-pre step (Nowak & Pucker, 2025).

Basecalling of sequencing data generated with R10.4.1 flow cells and stored in POD5 files was performed with dorado v0.8.3 (Oxford Nanopore Technologies, 2024).

### Direct RNA sequencing

RNA for direct RNA sequencing was extracted from one white petal with the Monarch Total RNA miniprep Kit (New England BioLabs) combined with the manufacturer specific DNase treatment. Homogenization was performed using the Bertin Ribolyser at 6500 RPM with two cycles of 30 seconds each, with a 10 second break between cycles. The extracted RNA quality and quantity was determined by NanoDrop measurement and additional Qubit measurement before conducting sequencing. Direct RNA sequencing was performed on a MinION Mk1B using an R9.4.1 flow cell and ONT’s direct RNA sequencing kit SQK-RNA002 following the suppliers’ protocol. Sequencing data were collected in FAST5 files and subjected to basecalling using Guppy v6.4.6+ae70e8f with the rna_r9.4.1_70bps_hac model and a minimum quality score of 7. Reads generated by direct RNA sequencing were aligned to the *V. cruziana* genome sequence (VB02) with minimap2 (v2.17 r941) using the following parameters: -ax splice -uf -k14 (Li, 2018). Sorting of the resulting SAM file and conversion into BAM was conducted with samtools v1.10 (Li *et al*., 2009). Mappings to the genome sequence were visualized in the context of the structural annotation with Integrative Genomics Viewer (IGV) v2.17.4 (Robinson *et al*., 2011). Direct RNA sequencing data was utilized to validate the structure of genes of interest through manual inspection of the long read alignments.

### Genome sequence assembly

An initial genome sequence assembly (VB01) was generated with all ONT reads using Shasta v0.10.0 (Shafin *et al*., 2020) with the config set to ‘Nanopore-May2022’. A second assembly (VB02) was generated with NextDenovo2 v2.5.2 (Hu *et al*., 2024) using the config file that is presented in Additional file 0. Whitespaces in contig names were removed with “sed ’s,,_*,g’ asm.fasta > asm.rm-wht-spc.fasta”.* General assembly statistics like assembly size, number of contigs, and N50 were calculated with contig_stats3.py (Meckoni *et al*., 2023). A completeness assessment was conducted with BUSCO v5.7.1 (Simão *et al*., 2015; Manni *et al*., 2021) using the genome or transcriptome mode according to the input data type and the reference dataset embryophyta_odb10. A third assembly (VB03) was generated with Verrko2 (v2.2.1) (Antipov *et al*., 2024) and scaffolded with CPhasing (v0.2.5) (https://wangyibin.github.io/CPhasing/latest/). Verrko2 was run solely with the HERRO corrected R10 reads and additional parameters: --par-run 13 464 48 --local-memory 464 --local-cpus 13. CPhasing input was the assembly resulting from Verkko2 and the Pore-C reads with additional parameters: -t 24 -n 12:1 -hcr -p CATG. Final assembly was then obtained by using agptools (bio_agptools, Version: 0.0.3, https://github.com/WarrenLab/agptools) to obtain a FASTA including the chromosome representing scaffolds and seqkit head -n 12 (v 2.9.0) (Shen *et al*., 2024) to filter it to include only the chromosome representing scaffolds.

### Structural annotation

The prediction of gene models was conducted with AUGUSTUS v3.3 (Stanke *et al*., 2006; Keller *et al*., 2011) for the assembly VB01 with parameters previously optimized for the detection of non-canonical splice sites (Pucker *et al*., 2017) (annotation VB01.v001). For the assembly VB02, BRAKER3 v3.0.8 (Gabriel *et al*., 2023) was applied with protein hints derived from Viridiplantae.fa (Kuznetsov *et al*., 2023) and RNA-seq hints. The RNA-seq hints were derived from a mapping of paired-end RNA-seq data (ERR12708812, ERR12708813, ERR12708814, ERR12708815, ERR12708816, ERR12708817, ERR12708818, ERR12708819, ERR12708820) generated with HISAT2 v2.2.1 (Kim *et al*., 2019) with default parameters against the assembly VB02. Samtools v1.10 (using htslib 1.10.2-3ubuntu0.1) (Li *et al*., 2009) was applied to generate a sorted BAM file which was then passed to BRAKER3. This resulted in the annotation VB02.v001. Annotation VB02.v002 was largely based on VB02.v001, only complemented with the coding sequence of an anthocyanin synthase (ANS) gene model lifted from the VB01 annotation. The derived polypeptide sequence of ANS was compared against the VB02 assembly via tBLASTn v2.15.0+ (Gertz et al., 2006) to locate this gene. Similarly like VB02.v001, VB02.v003 was partly generated with BRAKER3, but the Viridiplantae.fa protein hints have been combined with all proteins found on UniProt (The UniProt Consortium, 2021) for the Nymphaeaceae family as of 2024-09-02 (taxon id 4410, for all accession numbers see table Additional file 1a, https://doi.org/10.24355/dbbs.084-202406040628-0). The VB02.v003 BRAKER3 results have been merged with another annotation conducted with GeMoMa-1.9 (Keilwagen *et al*., 2016, 2019) running the GeMoMa pipeline with the same sorted BAM file and additional reference hints (*Nymphaea colorata* (GCF_008831285.2), *Nymphaea thermarum* (GCA_011799765.1), *Brasenia schreberi* (GCA_030142015), *Brasenia schreberi* 2 (GCA_036023115.1), *Amborella trichopoda* (GCA_000471905.1)) for which the parameter ‘s=own’ was specified. Additional parameters: ‘pc=true pgr=true o=true Extractor.r=true GAF.f=“start==’M’ and stop==’*’ and (isNaN(score) or score/aa>=’0.75’)” AnnotationFinalizer.r=NO’. For the combination with BRAKER3 results, the RNA-seq evidence was extracted with the GeMoMa tool ERE and annotation evidence from the BRAKER3 run was obtained by running GeMoMa tool AnnotationEvidence with the parameter ‘c=UNSTRANDED’, the BRAKER3 result GTF file and the extracted RNA-seq evidence. The resulting GFF file was used together with the GFF files generated for each reference species from the GeMoMaPipeline run with the GeMoMa tool GAF and additional parameters ‘f=“start==’M’ and stop==’*’ and aa>=15 and ((isNaN(score) and avgCov>0) or score/aa>=’3.50’)” atf=“evidence>1”’, to generate the filtered annotation. On this combined and filtered GFF file, GeMoMa tool AnnotationFinalizer with the options ‘tf=true rename=SIMPLE p=VB02 n=false’ and sub further GeMoMa tool Extractor with the options ‘p=true c=true identical=true’ was applied to generate cds and pep FASTA files (final annotation VB02.v003). VB03 structural annotation (VB03.v001) was generated like VB02.v003 with the following changes: Additional RNA-seq hints have been used (ERR14153978, ERR14153979, ERR14153980, ERR14153981, ERR14153982, ERR14153983, ERR14153984, ERR14153985, ERR14153986, ERR14153987, ERR14153988, ERR14153989, ERR14153990, ERR14153991, ERR14153992, ERR14153993, ERR14153994, ERR14153995, ERR15076234, ERR15076235, ERR15076236, ERR15076237, ERR15076238, ERR15076239, ERR15076240, ERR15076241, ERR15076242, SRR10158663), the additional protein hints from UniProt have been readjusted to contain all entries for *Nymphaeales* as of 2025-05-18 (taxon ID 261007, for all accession numbers see table Additional file 1b), an additional reference species for GeMoMa was used (GCA_000001735.2), the BRAKER3 annotation was filtered with GeMoMa GAF (parameters: f=“start==’M’ and stop==’*’ and aa>=15 and (isNaN(score) and avgCov>0)”) before merging it with all unfiltered predictions from the reference species with GeMoMa GAF (parameters: f=“start==’M’ and stop==’*’ and aa>=15 and (isNaN(score) or score/aa>=’3.25’)” atf=“tie==1 or sumWeight>6”), GeMoMa AnnotationFinalizer was run with the additional option “p=Vcurz_”, the resulting GFF3 file was filtered with AGAT (v1.4.1, agat_sp_fix_features_locations_duplicated.pl, additional parameter: -v) (https://github.com/NBISweden/AGAT).

### Functional annotation and candidate gene identification

Genes associated with the anthocyanin metabolism were identified with three dedicated tools. To identify the structural genes of the flavonoid biosynthesis pathway, an analysis with KIPEs v3.2.4 (Rempel *et al*., 2023) and the flavonoid baits data set v.3.1.7 was conducted (Additional file 2). Flavonoid biosynthesis controlling MYB transcription factors were annotated using the MYB_annotator v1.0.1 (Pucker, 2022) with parameters described in Additional file 3. The bHLH transcription factors were identified with the bHLH_annotator v1.04 (Thoben & Pucker, 2023) with parameters described in Additional file 4. A general annotation was produced using construct_anno.py and the functional annotation available for *A. thaliana* (Pucker & Iorizzo, 2023).

### RNA-seq data analysis

Quantification of transcript abundances (referred to as ‘gene expression’ in the following) was performed using kallisto v0.48.0 (Bray *et al*., 2016) on the predicted coding sequence. Count tables were merged using a customized python script as well as gene count normalization by transcript per million (TPM) (Pucker & Iorizzo, 2023). These results provided the basis for visualization of gene expression for the structural genes and transcription factors in heat maps. Principal component analysis (PCA) using ggplot2 v3.4.4 library was performed on transcripts with counts >3. For initial analysis of transcriptome data, DESeq2 v1.42.0 (Love *et al*., 2014) using BiocManager v1.30.22 and matrixStats v1.2.0 was executed in R v2023.09.1+494. Gene candidates were determined through the described functional annotation and by sequence alignment using DIAMOND v2.1.9 (Buchfink *et al*., 2015, 2021). Visualization of final gene candidates with predicted annotation and z-score normalization was conducted using customized python scripts (https://github.com/bpucker/vici) based on seaborn v0.13.2 (Waskom, 2021).

### Phylogenetic analysis

For phylogenetic classifications, sequences of *V. cruziana* MYB candidates in subgroup 4, 5, 6 and 7 were aligned with the *A. thaliana* bait sequences of the MYB_annotator (Pucker, 2022) and KcMYB1 (Huang *et al*., 2022) via MAFFT v7.526 (Katoh & Standley, 2013). IQ-TREE v2.0.7 (Nguyen *et al*., 2015; Minh *et al*., 2020) was applied to infer a maximum likelihood tree with the parameters -alrt 1000 -B 1000. The final tree file was visualized in iTOL (Letunic & Bork, 2021).

### Investigation of environmental factors

To investigate whether pollinators have a direct impact on anthocyanin biosynthesis, young flower buds of *V. cruziana* were wrapped up in Nylon nets to ensure that no pollinators can enter the flower. After completion of the flowering period, the pigmentation status was documented. Furthermore, to determine if light exposure could be the decisive factor for anthocyanin production in the petals, flower buds have been covered with aluminum foil directly after they emerged. Several flowers were unwrapped and investigated regarding their pigmentation after the flowering period was completed. In some other cases, the aluminum foil was partially removed at the time point where the fruity smell occurred. This approach was designed to show whether there are selective differences in the pigmentation status after sunlight exposure.

Additionally, the development of several flowers was monitored on a daily basis to determine the development timeframe from bud to a fully opened flower. The morphological characteristics and timespan of floral growth of emerging flowers have been captured whereby a variation has been used for statistical analysis. We analyzed the diameters from bud to blossom as well as the average duration for flower development in respect to environmental influences.

## Results

### Assembly results

A completeness assessment revealed 92.4%,93.6%, and 95.1% complete BUSCO genes for the VB01 (Shasta),VB02 (NextDenovo2), and VB03 (Verkko2+CPhasing) assembly, respectively. VB02 has a size of 3.54 Gbp comprising 837 contigs with an N50 length of 14.3 Mbp. VB03 has a size of 3.50 Gbp comprising 12 scaffolds with an N50 length of 300 Mbp (VB03.genome.pore-c_contact_map.png, https://doi.org/10.60507/FK2/5DS0JZ). In total, 50,681 (VB02.v001), 50,682 (VB02.v002) and 25,035 (VB02.v003) protein coding genes were annotated in VB02 and 27158 (VB03.v001) in VB03. The assembled genome sequence and the corresponding annotations are available via LeoPARD (https://doi.org/10.24355/dbbs.084-202406040628-0 for VB02.v001 and VB02.v002 and https://doi.org/10.24355/dbbs.084-202502270747-0 for VB02.v003), bonndata (https://doi.org/10.60507/FK2/5DS0JZ for VB03.v001) and NCBI (GCA_965616825.1 also for VB03.v001).

### Structural genes of the anthocyanin biosynthesis are differentially expressed between white and light pink petals

A principal component analysis (PCA) of the transcriptomic data of young and mature petals revealed distinct clusters of the replicates derived from white and light pink flower samples, respectively, supporting clear differences between the two groups and homogeneity within each group (Additional file 5).

To understand the molecular basis of the flower color variations in *V. cruziana*, structural genes required for anthocyanin biosynthesis were identified. Expression profiles of the best candidates were analyzed with respect to the white and light pink flower color in two developmental stages (**Figure 4**). The first stage is referred to as young petals and represents buds between day 3 and day 8 whereby the second stage corresponds to mature petals (opened flower). In both stages, the majority of genes required for anthocyanin biosynthesis (*F3’H*, *F3’5’H*, *DFR*, *ANS*, *arGST*, and *UFGT*) exhibited higher expression levels in anthocyanin-pigmented light pink petals compared to white petals (Additional file 6). The expression pattern of *CHS* and *CHI* matches better to the flavonol biosynthesis gene *FLS* than to the anthocyanin biosynthesis gene *DFR*, i.e., *CHS* and *CHI* exhibited a higher expression within the mature white petals. However, in young petals *CHS* and *CHI* exhibit higher expression values within the light pink petals although the pattern does not completely align with *DFR* expression. To convert flavanones into dihydroflavonols, *F3H* is required and showed increased expression levels in the samples taken from light pink petals in both developmental stages. Branching reactions towards delphindin or cyanidin are catalyzed by F3’5’H and F3’H, respectively. The corresponding genes are present in *V. cruziana* whereby only light pink mature petals express *F3’5’H.* In total *F3’H* exhibited a higher activity in young light pink petals compared to white whereby a varying activity can be observed between coloration in mature petals. Dihydroflavonols are converted to the precursors of anthocyanidins, the leucoanthocyanidins, by DFR. Higher expression levels of *DFR* were observed in mature light pink petals compared to white petals whereas young petals had a similar tendency of higher *DFR* expression within light pink petals. The production of anthocyanidins requires also the downstream enzymes ANS and arGST to catalyze the conversion into the anthocyanidins delphinidin and cyanidin in *V. cruziana. ANS* is moderately more active in young white petals and conversely in mature petals. Nevertheless *arGST* expression is significantly increased in the light pink petals of young and mature petals (**Figure 4**).

**Figure 4:**
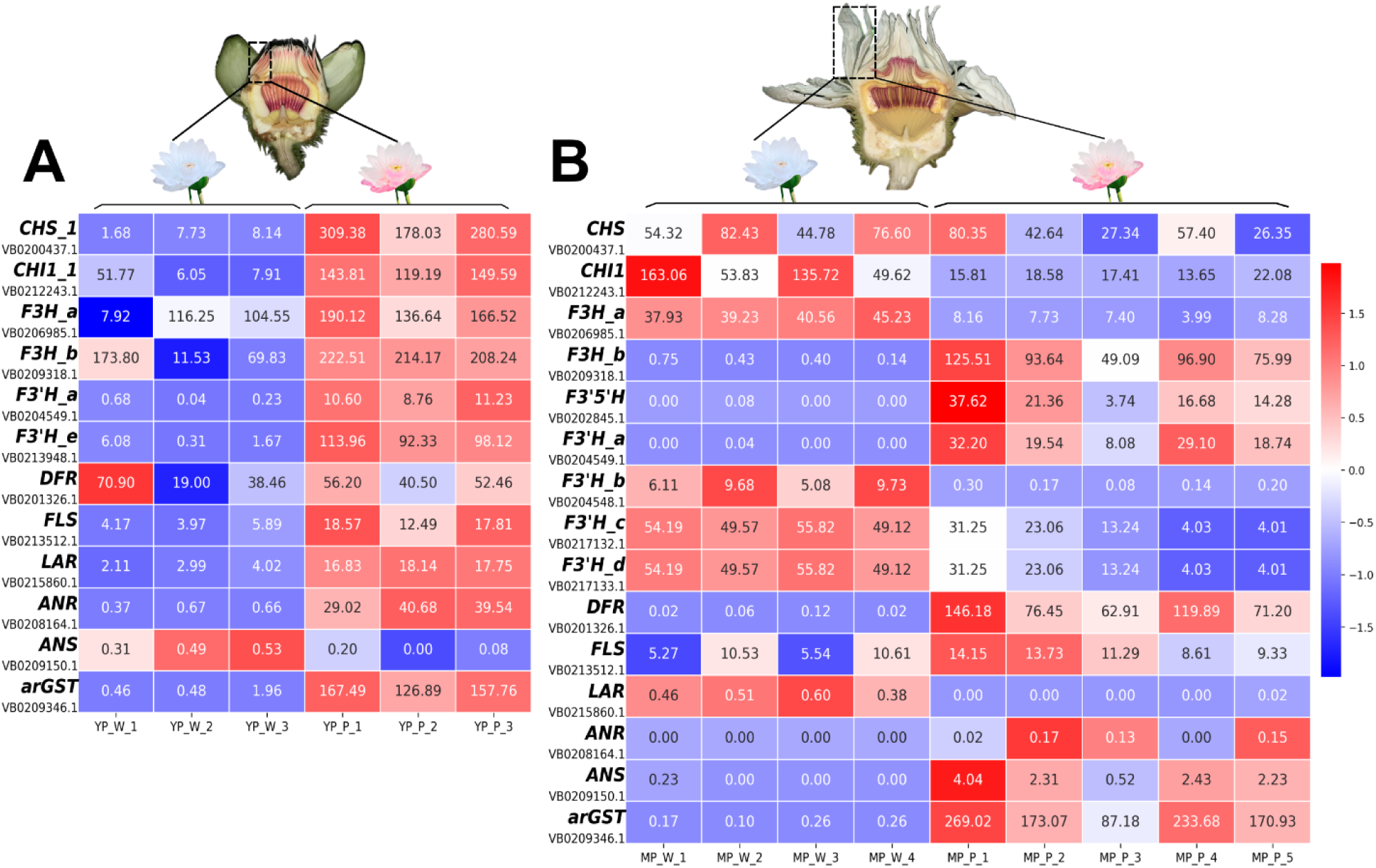
Expression pattern of structural genes associated with the flavonoid biosynthesis in young (A) and mature (B) *V. cruziana* flowers, each compared between white and light pink petals. *PAL* (*Phenylalanine ammonia lyase*), *C4H* (*Cinnamate 4-hydroxylase*), *4CL* (*4-coumarate-CoA ligase*), *CHS* (*Chalcone synthase*), *CHI* (*Chalcone isomerase*), *F3H* (*Flavanone 3-hydroxylase*), *F3’H* (*Flavonoid 3’-hydroxylase*), *F3’5’H* (*Flavonoid 3’,5’-hydroxylase*), *FLS* (*Flavonol synthase*), *DFR* (*Dihydroflavonol 4-reductase*), *LAR* (*Leucoanthocyanidin reductase*), *ANR* (*Anthocyanidin reductase*), *ANS* (*Anthocyanidin synthase*), *arGST* (*anthocyanin-related Glutathione S-transferase*). Expression values are z-score normalized TPM values. Values given in the individual fields of the heatmap are TPM values. YP: Young Petals, MP: Mature Petals; W: white, P: light pink; 1-5: Indicating replicates.

### Expression of anthocyanin biosynthesis regulators differs between white and light pinkish flowers

Anthocyanin biosynthesis is regulated by a complex comprising transcription factors from the MYB family, the bHLH family, and a specific WD40 protein (Gonzalez *et al*., 2008). After identification of the required structural genes, best candidates for potential transcriptional regulators were analyzed. MYBs belonging to subgroup 6 (*VcrMYB-SG6*) are orthologous to previously described activators of the anthocyanin biosynthesis. Two candidates have been identified as active in young petals, although they appear not to be expressed in mature petals (**Figure 5**). As one transcription factor with activatory properties for proanthocyanidin production, one candidate for subgroup 5 MYBs (*VcrMYB123*) was identified. In young and mature petals, *VcrMYB123* tends to be more active in light pink petals compared to white petals. Flavonol production is mainly regulated through MYBs of subgroup 7 (Stracke *et al*., 2007, 2010). Expression patterns of corresponding gene candidates (*VcrMYB-SG7*) exhibit activity in light pink petals of young petals whereas no transcripts have been detected in mature petals. Additionally, *VcrTT8* encoding a bHLH protein and *VcrTTG1* encoding the WD40 protein of the MBW complex, were predicted in the functional annotation process. *VcrTT8* tends to show higher expression in light pink petals at both young and mature petals. The corresponding gene candidate for *VcrTTG1* does not exhibit clear expression differences between white and light pink young petals, whereas increased activity is observed in light pink mature petals. Further, *VcrMYB-SG4* candidates were identified, which could be orthologues of the anthocyanin biosynthesis repressing regulator *AtMYB004*. Expression levels of *VcrMYB004_a* are increased within the white mature petals thus showing a pattern opposite to anthocyanin accumulation. However, *VcrMYB004_a* in young petals and another potential *AtMYB004* ortholog, *VcrMYB004_b* exhibited higher transcript abundance in light pink petals compared to white (**Figure 5**).

**Figure 5:**
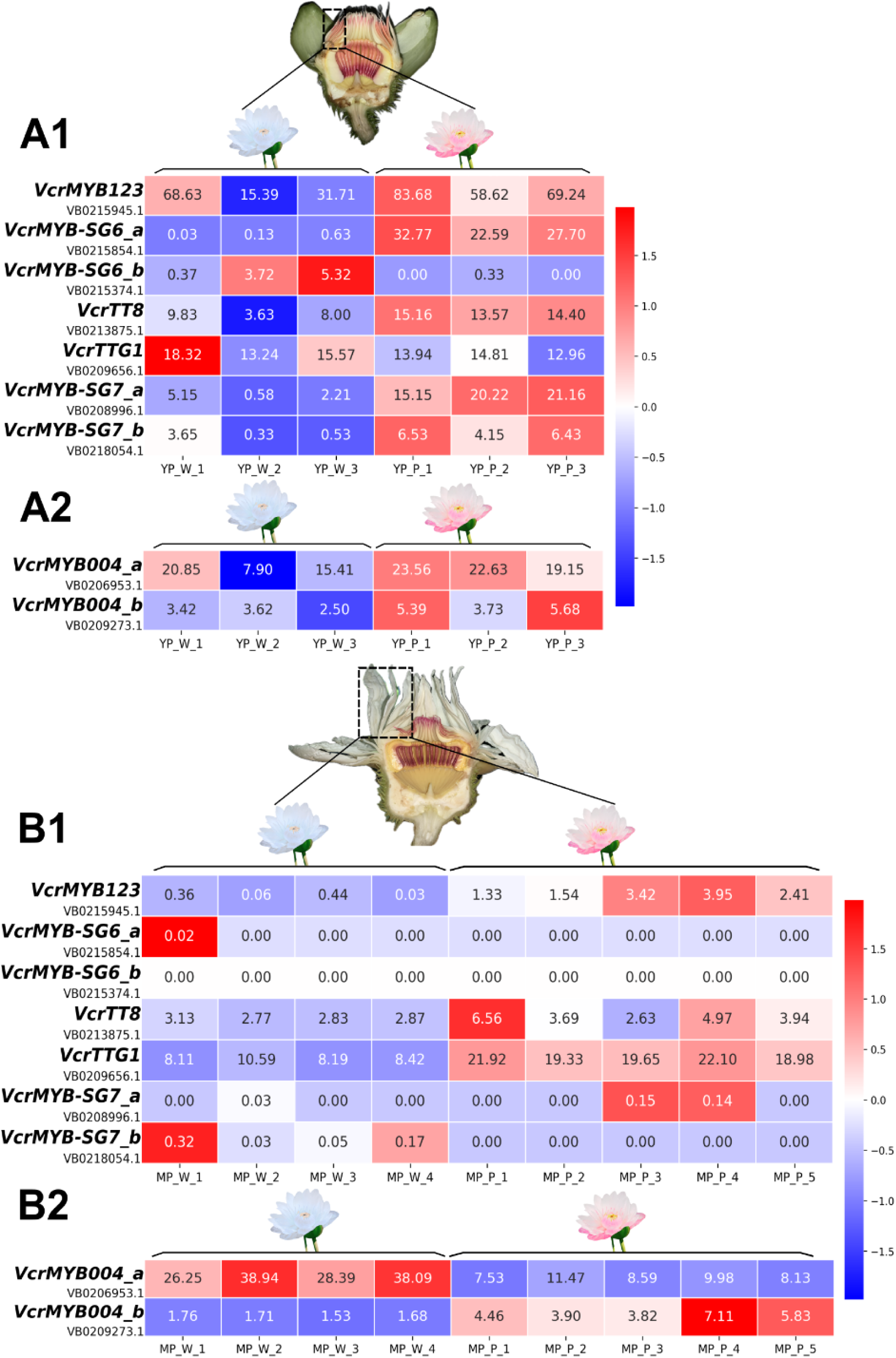
A1-B1: Expression pattern of transcription factor genes putatively required for anthocyanin biosynthesis activation in young (A1) and mature (B1) petals of *V. cruziana*. *VcrMYB123* belongs to subgroup 5 MYBs required for proanthocyanidin production whereas subgroup 6 MYBs activate the anthocyanin biosynthesis. *VcrMYB123* and and *VcrMYB-SG6* expression patterns align partially with the anthocyanin accumulation in young petals whereby *VcrMYB123* exclusively seems to regulate anthocyanin production in mature petals. *VcrTT8* represents the associated bHLH transcription factor and *VcrTTG1* edcodes a WD40 protein required for anthocyanin biosynthesis activation. A2-B2: *VcrMYB004_a* and *VcrMYB004_b* probably possess flavonoid biosynthesis repressing activities as they belong to the subgroup 4 MYBs. Numeration of annotated gene IDs indicates affiliation to the same functionally annotated candidate. Expression values are z-score normalized TPM values. If a candidate gene shows zero TPM, the z-score has been assigned a value of zero. Values given in the individual fields of the heatmap are TPM values. YP: Young Petals, MP: Mature Petals; W: white, P: light pink; 1-5: Indicating replicates.

To explore other MYB transcription factors that have been reported in the context of anthocyanin biosynthesis in species closely related to *V.cruiziana*, a search for the ortholog of the recently published KcMYB1 from *Kadsura coccinea* was conducted. KcMYB1 belongs to the MYB subgroup 6 and overexpression in *Nicotiana benthamiana* resulted in anthocyanin accumulation (Huang *et al*., 2022). Using functional annotation and a phylogenetic analysis, VB0215945.1 was classified as a subgroup 5 R2R3-MYB, while KcMYB1 was located in the subgroup 6 clade close to the *V. cruziana* genes VB0215854.1, VB0215374.1, and VB0215373.1 (Additional file 8). However, only VB0215854.1 and VB0215374.1 (*VcrMYB-SG6*) were substantially supported by aligned RNA-seq reads, indicating a promising SG6 MYB in *V. cruziana*. In contrast, genes without RNA-seq support might be pseudogenes and have not been picked up as differentially expressed between white and light pinkish petal samples in both stages (Additional file 8, Additional file 6, Additional file 7).

### Sunlight appears as a decisive factor in flower color change

Sunlight is a well studied inducer of anthocyanin pigment (Albert *et al*., 2009; Araguirang & Richter, 2022; Grünig *et al*., 2024). To explore the contribution of sunlight and pollinators, a range of experiments was conducted. Some flowering plants induce flower color changes as an indication for pollination. These visible changes serve as a communication signal between the plant and their pollinator. Despite the shielding from pollinators, light pink flowers were observed after completion of the flowering period. This indicates that pollinators are not the driving force of the anthocyanin accumulation in flowers. However, it is also known that anthocyanin biosynthesis can be induced upon light exposure. To confirm whether light is the cause for flower color change, emerging flower buds have been wrapped up in aluminum foil and later investigated in respect to their pigmentation status. Completely covered flower buds (i.e. without any exposure to sunlight) exhibited no significant pigmentation after their flower period ended. Albeit, the flowers where aluminum foil was partially removed showed selective pigmentation in the exposed areas compared to those areas that remained covered (**Figure 6**). These findings indicate that light exposure might play an essential role in the anthocyanin production in flower coloration of *V. cruziana*.

**Figure 6:**
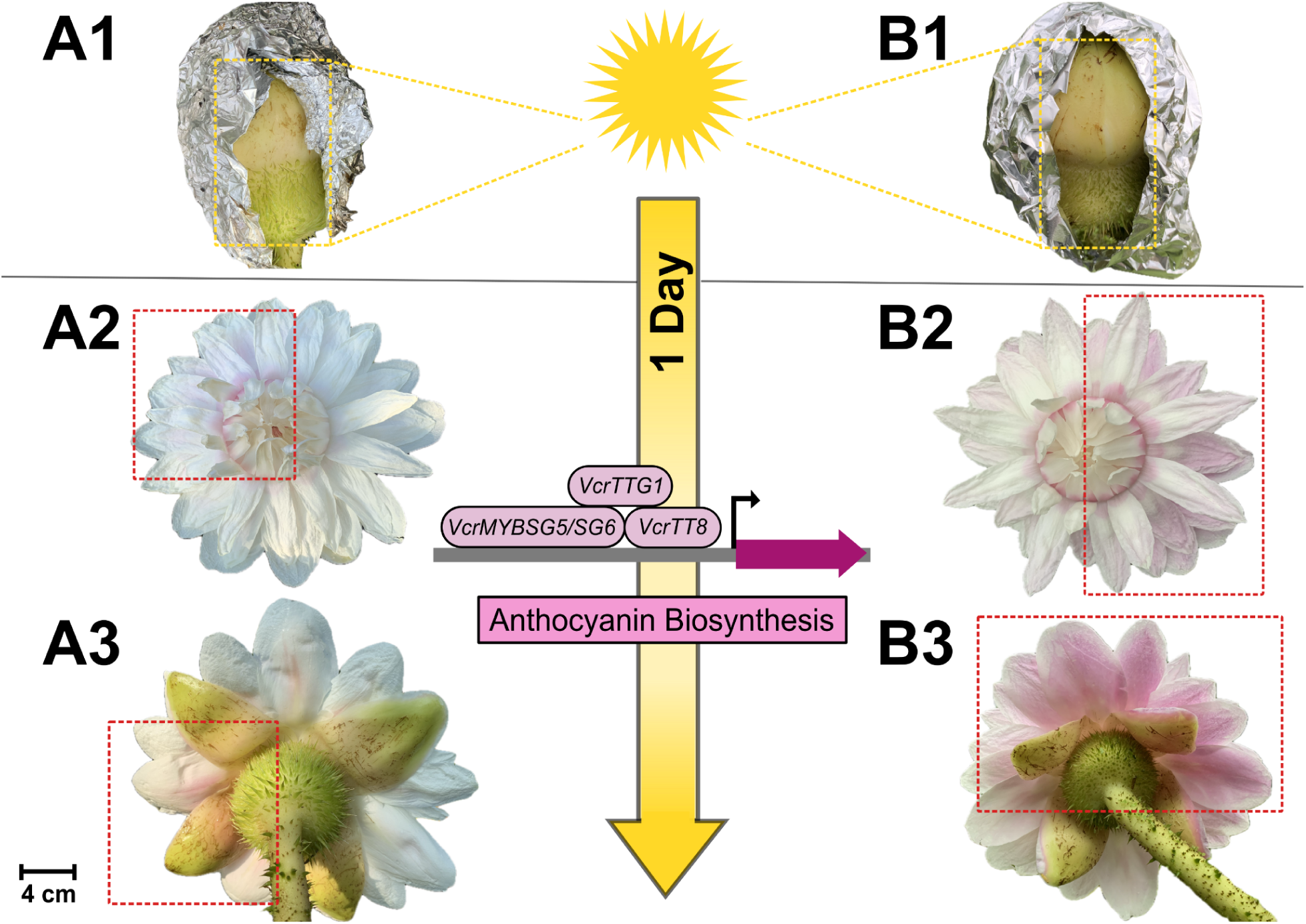
Illustration of sunlight exposure experimental workflow including representative results. Buds have been completely covered in aluminum foil as they emerged to prevent sunlight exposure. A1-B1: Aluminum foil was later partially removed when the fruity smell occurred. The yellow dotted line represents the area of sunlight exposure. A2-B2: Front side of the fully opened flower 24h after partial removal of the foil. A3-B3: Backside of the fully opened flower 24h after partial removal of the foil. Red dotted lines are representing the area of sunlight exposure on the fully open flower. Distinct coloration patterns are observed when comparing sunlight exposed regions to those that remain unexposed. Scale Bar: 4 cm.

### Flower development time of *V. cruziana* depends on environmental conditions

Flower development is a fundamental process in the life cycle of plants. Timing of flowering and growth dynamics differ across species. Flower size and morphologies change dynamically during the growing processes, reflecting both intrinsic programs and external influences such as temperature. **Figure 7** shows exemplarily the development from bud to a fully open flower within 15 days.

**Figure 7:**
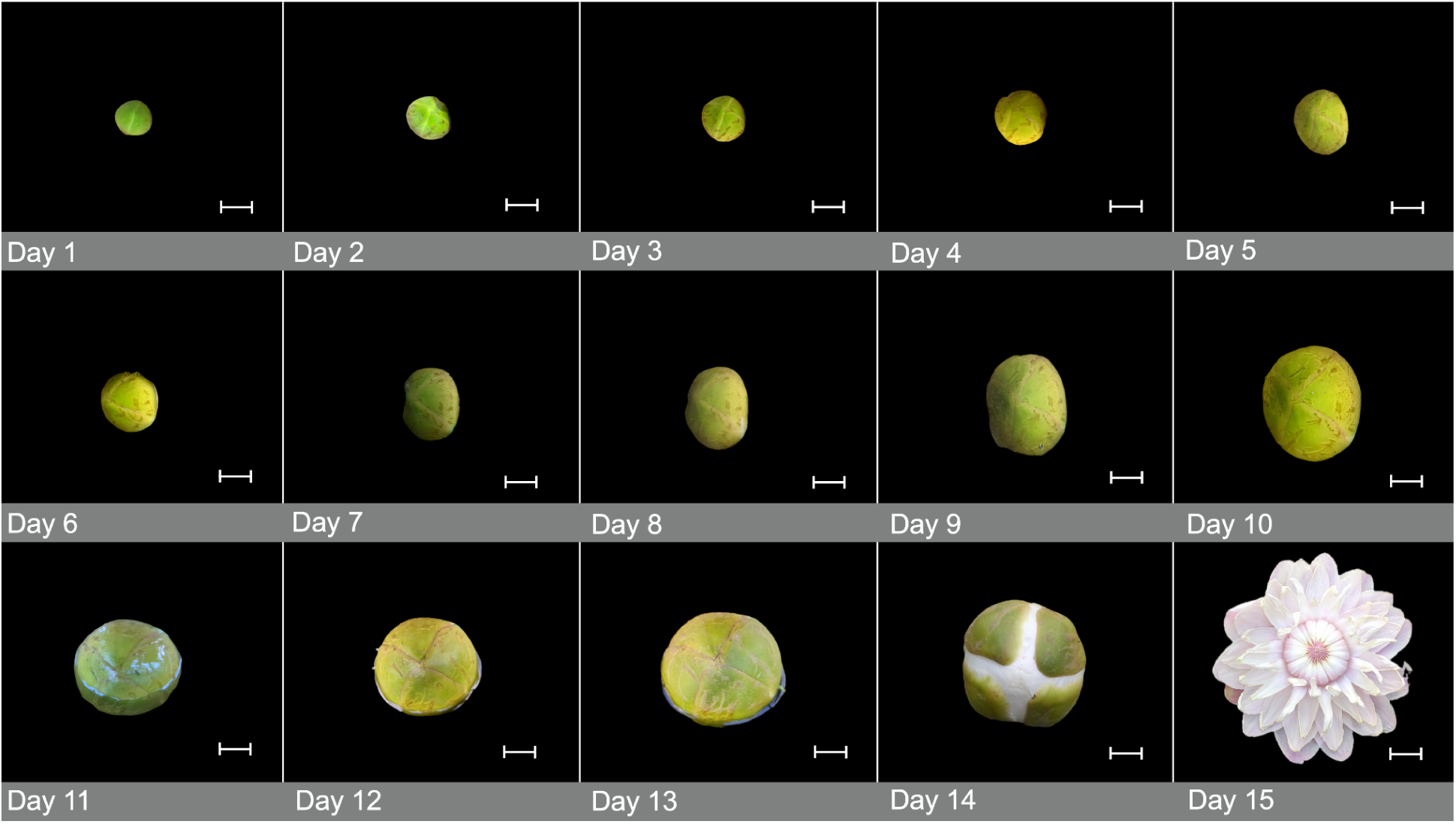
Exemplary representation of bud development of *V. cruziana* to a fully open flower. Scale bar: 2.5 cm

To quantify the development from bud to blossom, we measured the diameter of five selected buds, including the opened flower within a timeframe of 15 days. Green violin plots are representing bud diameters whereas light pink shows the diameters of the fully opened flowers. Opened flowers reach span widths between 15 cm and 25 cm (**Figure 8**).

**Figure 8:**
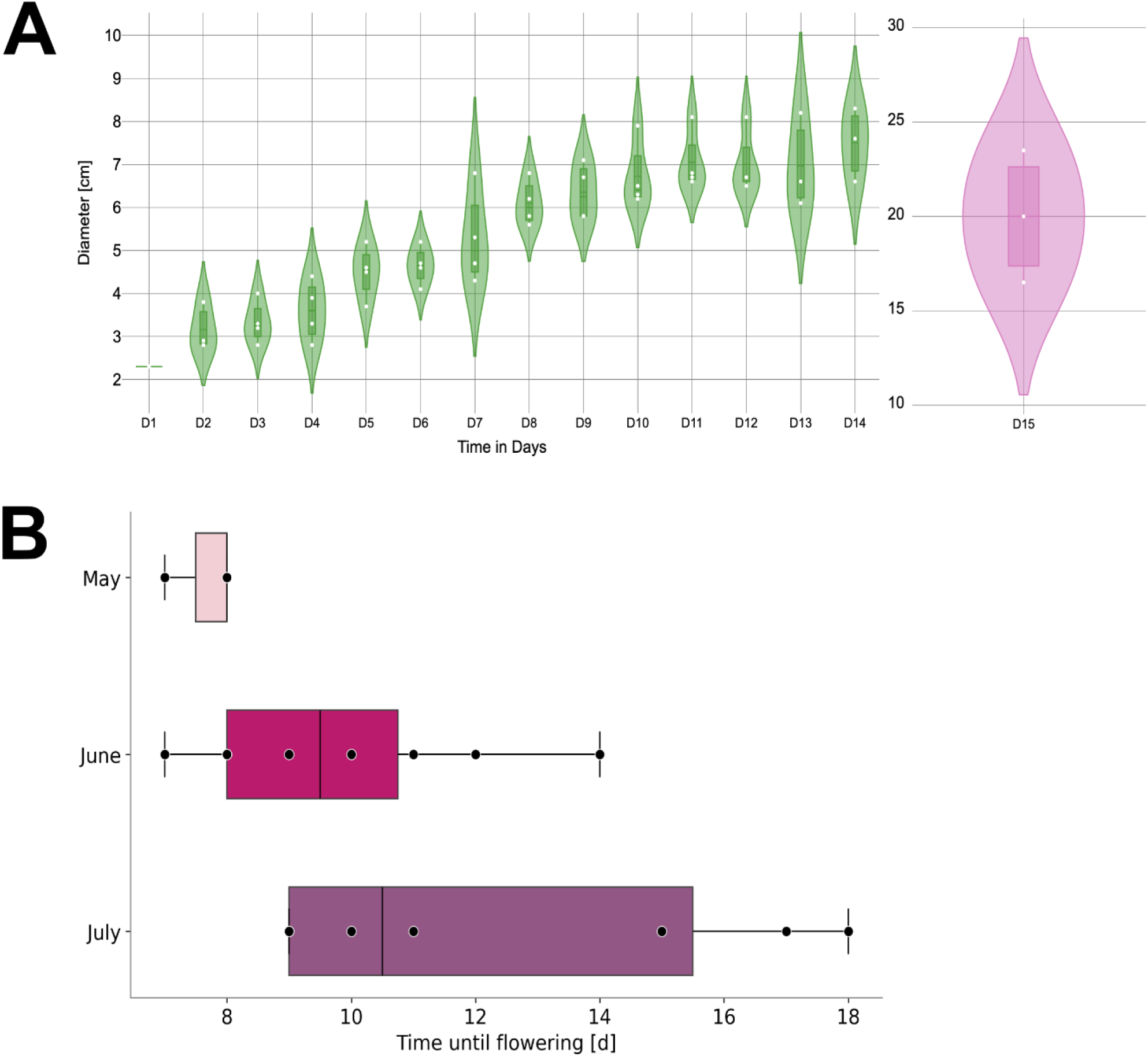
Developmental aspects of *V. cruziana*. Violin plot showing the diameter of buds during development (green) and fully opened flower (light pink). Measurements of buds have been conducted in a time frame of 15 days. n: 5 (A). Boxplots illustrates the average development time for each month with a scatter overlay representing the duration of individual bud to flower. Documentation started as soon as bud appeared. May n:3; June n:10; July n:8. Median is indicated by the central line within the boxes (B).

Noticeable is that the developmental period of *V. cruziana* flowers differs from bud to bud and might be influenced by the time of the year. The associated observations are shown in **Figure 8**, representing the average developmental time for each month combined with the duration of individual bud to flower timespans. It appears that the flowering period is extended in the later months as timeframes range from 7 to 8 days in May, 7 to 14 in June, and 9 to 18 in July.

## Discussion

### Differential expression of flavonoid biosynthesis genes explains anthocyanin production in light pink petals of *V. cruziana*

The contiguous genome sequence of *V. cruziana* and the corresponding annotation enabled the identification of genes encoding relevant proteins and transcription factors required for the anthocyanin biosynthesis (**Figure 6**). The differential expression patterns compared between white and light pink petals align with the active anthocyanin production in the light pink flower.

The first committed genes of the flavonoid biosynthesis (*CHS*, *CHI*) were higher expressed in the young white flower likely initiating the start of anthocyanin production. Additionally, *FLS* and *LAR* expression patterns did not match the expression level patterns of the other genes in mature petals. The *FLS* pattern can be due to its activity in flavonol biosynthesis which is active in the white petals, supported by Wu *et al.,* 2018 who identified flavonols in white *V. cruziana* flowers. Furthermore, *F3’5’H* as the branching point towards delphinidin production exhibited increased expression levels in mature light pink petals, whereas no transcripts have been detected in young petals. This finding is surprising as delphinidin is one anthocyanidin required for blue flower colors which has not been observed in *Victoria spp*. flowers. However, delphinidin derivatives were identified besides cyanidin derivatives as the major anthocyanins in the *V. cruziana* and *V. amazonica* (Wu *et al*., 2018). Perhaps blue colors are not visible due to overlay with cyanidin derivatives. F3’H represents another branching point in the flavonoid biosynthesis branch that leads to cyanidin. However, multiple transcripts of different gene candidates have been observed in mature petals which may represent distinct transcript isoforms. Another possibility could be recently generated gene duplicates which cannot be precisly distinguished. Also, technical limitations within the functional annotation cannot be excluded as some apparently crucial amino acid residues might be missing due to the distant position of *V. cruziana* in the phylogenetic tree. As *ANS* and *arGST* encode major enzymes in anthocyanin biosynthesis, their expression pattern matches the anthocyanin accumulation in mature light pink petals and partially in light pink young petals. Interestingly the *arGST* expression is significantly higher in light pink petals of both developmental stages compared to the corresponding *ANS* expression. The substantially higher expression of *arGST* compared to *ANS* might ensure the channeling of metabolites towards anthocyanins. Previous studies identified arGST as a decisive step in the anthocyanin biosynthesis (Eichenberger *et al*., 2023; Pucker *et al*., 2024).

### *Vcr*MYB123 and *Vcr*MYB-SG6 may interact with *Vcr*TT8 and *Vcr*TTG1 to activate the anthocyanin biosynthesis in light pink flowers of *V. cruziana*

Transcription factors fulfill a crucial role when it comes to orchestrating gene expression. Essential for anthocyanin production in plants are MYB proteins, regulating various branches of the pathway. To understand why anthocyanin pigments are accumulating in flowers, the responsible transcription factor needs to be identified. *VcrMYB123*, most likely an *AtMYB123* orthologue, was identified in *V. cruziana. Vcr*MYB-SG6 candidates were identified as potential orthologues of prominent anthocyanin-regulating MYBs like *MYB75/MYB90/MYB113/MYB114* in *A. thaliana.* Furthermore, candidates were identified as potential members of the flavonol-regulating MYB subgroup 7. It appears likely that VcrMYB-SG6 and potentially VcrMYB123 are the main anthocyanin-regulating MYBs in *V. cruziana*. Other gene candidates proposed for subgroup 6 are not supported by petal-derived RNA-seq data which could indicate that they might be active in other plant parts. Previous studies reported a role of subgroup 5 MYB (MYB5, MYB123) in activating the anthocyanin biosynthesis (Martínez-Rivas *et al*., 2023; Jiang *et al*., 2023) thus corroborating the putative role of *VcrMYB123* in this context. The additional sequence alignment of *KcMYB1* TF (JHA, Additional file 7) from *K. coccinea* further supports the assumption that VB0215854.1 and VB0215374.1 gene candidates encode anthocyanin activating subgroup 6 MYB transcription factors as *K. coccinea* belongs to the *Schisandraceae* (Huang *et al*., 2022), a basal angiosperm lineage like the *Nymphaceaee*. Further, the *VcrMYB004_a* and *VcrMYB004_b*, potential orthologues of the repressor *AtMYB004*, showed different expression levels in both stages. *VcrMYB004_a* shows partially contrasting transcript abundances as they are higher in white mature petals compared to light pink. In the young stage, *VcrMYB004_a* tends to be higher expressed in light pink petals compared to white. For *VcrMYB004_a in* one replicate, transcript levels in young white petals are similar to the light pink young petals. This could suggest a repression on anthocyanin production already setting in as the samples have been harvested from buds between day 3 and day 8. However, *VcrMYB004_b* exhibits higher expression levels in the young and mature light pink compared to the white flower, suggesting a repressive function on anthocyanin production in the light pink petals. This can be further supported by a study, which reported another AtMYB004 orthologue in *Musa* spp., *MaMYB4*, with repressive function on the anthocyanin biosynthesis genes (Deng *et al*., 2021). Li et al. observed decreased transcript levels in *CsMYB4a* transgenic plants suggesting a repressive function on the flavonoid biosynthesis pathway (Li *et al*., 2017).

**Figure 9:**
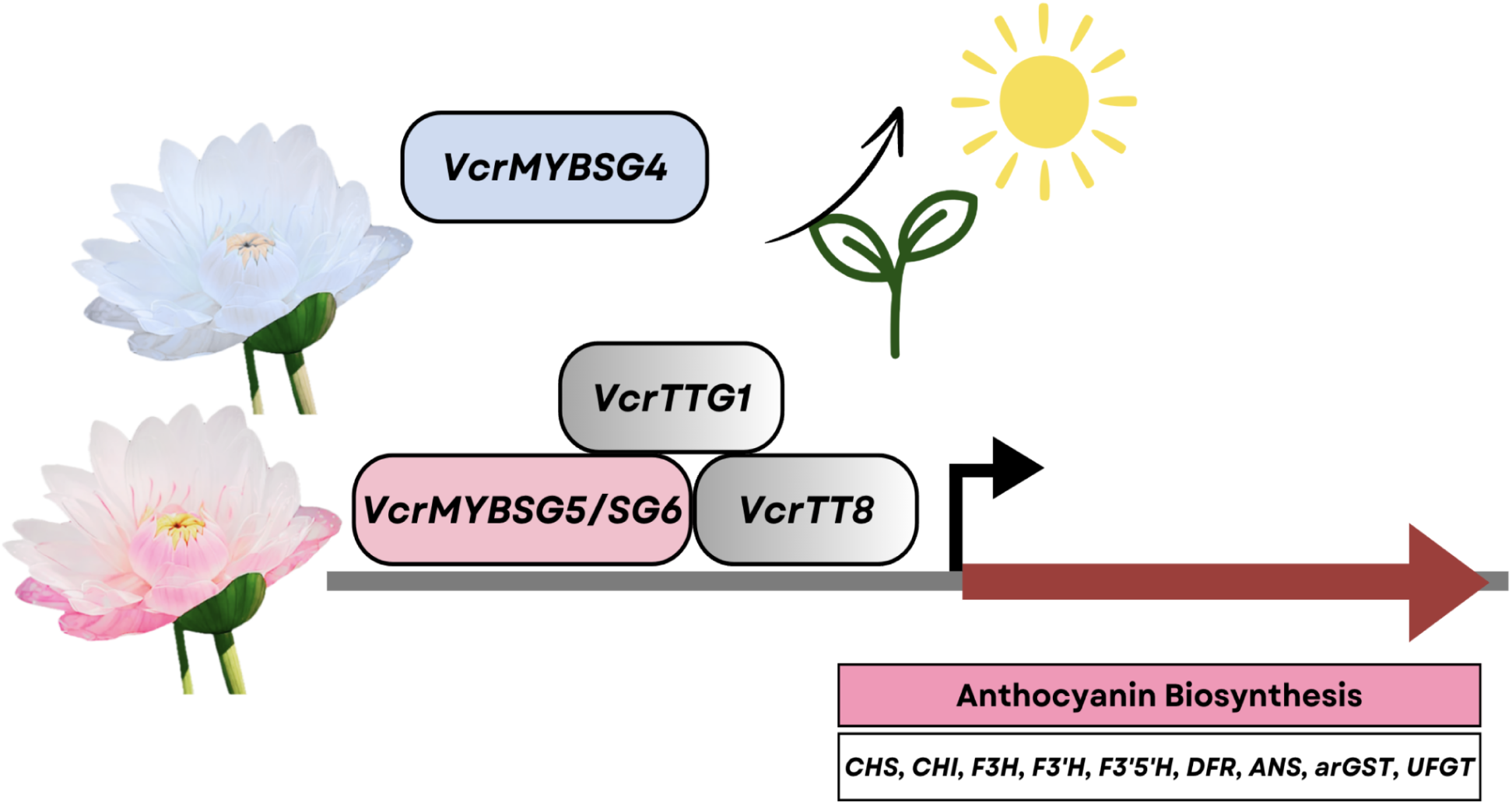
Graphical abstract summarizing the molecular mechanism involved in anthocyanin biosynthesis and potential factors triggering the flower color change of *V.cruziana. VcrMYB-SG4* gene candidates are partially active in white petals whereas additional activities are observed in light pink samples. *VcrMYB123* and *VcrMYB-SG6* are expressed in light pink petals within both stages. Factors that could trigger anthocyanin biosynthesis, thereby activating TFs and structural genes, might be light exposure or a general developmental program of the plant.

Based on a continuous genome sequence, we successfully identified all structural genes required for anthocyanidin production in *V. cruziana*. However, there is potential to improve the structural annotation as 93.6% complete BUSCOs have been discovered. The inclusion of additional RNA-seq hints derived from different plant parts might contribute to further improvements of the structural gene prediction and subsequently result in an improved functional annotation.

Further, we identified members of the *VcrMYB-SG6* and *VcrMYB123* as potential transcriptional activators of anthocyanin and proanthocyanidin biosynthesis based on correlation of their expression patterns with anthocyanin accumulation and phylogenetic analyses. To confirm the anthocyanin biosynthesis activation properties, the genes could be introduced into *A. thaliana myb75* mutants to conduct a complementation experiment. Besides, expression levels of the structural genes as well as transcription factors of *V. cruziana* could be analyzed in leaves and other parts of the plant to further investigate potential TFs activating structural genes in the flavonoid biosynthesis.

To reveal developmental and environmental factors triggering the coloration of *V. cruziana* flowers, we performed experiments to control light exposure and pollinator access to the flowers. Since *V. cruziana* is only opening its flowers at night, light triggering the anthocyanin formation inside the flower would have to pass through the outer petals. Potentially the first sunlight of the next day could be sufficient to induce anthocyanin biosynthesis in the outer petals of mature flowers. Further experiments are needed to fully understand the ecological relevance and environmental factors triggering the floral color transition through anthocyanin formation in *V. cruziana*.

## Declarations

### Ethics approval and consent to participate

Not applicable.

### Consent for publication

Not applicable.

### Data availability

Genomic sequencing data (PRJEB63973), direct RNA sequencing data (PRJEB63453), and RNA-seq data (PRJEB63973) are available via the European Nucleotide Archive (ENA). The assembled *Victoria cruziana* genome sequence VB02 and the corresponding annotation are available via LeoPARD (https://doi.org/10.24355/dbbs.084-202502270747-0). The genome sequence VB03 and the corresponding annotation VB03.v001 are available via bonndata (https://doi.org/10.60507/FK2/5DS0JZ) and NCBI (GCA_965616825.1).

### Authors’ contributions

MSN, BH, SNM, BP planned the study. MSN, BH, RF, KW, and BP performed sampling, nucleic acid extraction, and ONT sequencing. MSN, SNM, BH, KW and BP conducted bioinformatic analyses. MSN, BH, SNM, and BP interpreted the results and wrote the manuscript. All authors have read the final version of the manuscript and approved its submission.

## Acknowledgements

We thank Thorsten Marschall, Michael Kraft, and the entire team operating the Botanical Garden of TU Braunschweig for their excellent technical support. This work was supported by the BMBF-funded de.NBI Cloud within the German Network for Bioinformatics Infrastructure (de.NBI) (031A532B, 031A533A, 031A533B, 031A534A, 031A535A, 031A537A, 031A537B, 031A537C, 031A537D, 031A538A). We also thank all students participating in our ‘Data Literacy in Genome Research’ course and members of the research group Plant Biotechnology and Bioinformatics for discussion and support.

## Supplementary

**Additional file 0:** NextDenovo2 config file used for the assembly.

**Additional file 1a**: UniProt IDs of *Nymphaeaceae* family (taxon id 4410) reference proteins used for structural gene annotation (as of 2024-09-02).

**Additional file 1b**: UniProt IDs of *Nymphaeales* (taxon ID 261007) reference proteins used for structural gene annotation (as of 2025-05-18).

**Additional file 2:** Documentation of the KIPEs3 analysis.

**Additional file 3:** Documentation of the MYB_annotator analysis.

**Additional file 4:** Documentation of the bHLH_annotator analysis.

**Additional file 5:** Principal Component Analysis of the RNA-seq samples for young and mature petals confirming the clustering of biological replicates in their respective groups.

**Additional file 6:** Table containing the DESeq2 results of the white vs. light pinkish flower stage comparison in young inner petals.

**Additional file 7:** Table containing the DESeq2 results of the white vs. light pinkish flower stage comparison in Mature petals.

**Additional file 8:** Phylogenetic classification of VcrMYB candidates. Subgroup 6 (red), Subgroup 5 (brown), Subgroup 7 (yellow), Subgroup 4 (blue).

**Additional file 9:** Table containing TPM data.

**Additional file 10:** Table containing visualization of flowers after sunlight exposure experiment.

