## Additional file 0 for "Genome sequence and RNA-seq analysis reveal genetic basis of flower coloration in the giant water lily *Victoria cruziana*"

[General]
job_type = local # local, slurm, sge, pbs, lsf
job_prefix = nextDenovo_v2
task = all # all, correct, assemble
rewrite = yes # yes/no
deltmp = yes 
parallel_jobs = 20 # number of tasks used to run in parallel
input_type = raw # raw, corrected
read_type = ont # clr, ont, hifi
input_fofn = /vol/data/samuel/vici/v.cruziana_ND_assenmbyl_no_2/input.fofn
workdir = /vol/data/samuel/vici/v.cruziana_ND_assenmbyl_no_2/workdir

[correct_option]
read_cutoff = 5k
genome_size = 3300mg # estimated genome size
sort_options = -m 400g -t 10
minimap2_options_raw = -t 10
pa_correction = 10 # number of corrected tasks used to run in parallel, each corrected task requires ~TOTAL_INPUT_BASES/4 bytes of memory usage.
correction_options = -p 20

[assemble_option]
minimap2_options_cns = -t 10 
nextgraph_options = -a 1

# see https://nextdenovo.readthedocs.io/en/latest/OPTION.html for a detailed introduction about all the parameters
