## Additional file 3 for "Genome sequence and RNA-seq analysis reveal genetic basis of flower coloration in the giant water lily *Victoria cruziana*"

Please cite 'Pucker B (2022). Automatic identification and annotation of MYB gene family members in plants. BMC Genomics 23, 220 (2022). doi: 10.1186/s12864-022-08452-5' when using MYB_annotator.py.

MYB_annotator.py version: v1.0.1
MYB bait file: /xxx/MYB_annotator/MYB_baits.fasta	f92f35b0d351312c43a6e1b30aa0567e
MYB info file: /xxx/MYB_annotator/MYB_baits.txt	5836d7f542b672365883fa0a29d5d5a0
Subject FASTA file: /xxx/GeMoMa/6_combi_3_05_finalised_02/proteins.fasta	73d58054958954b1d221fcf612bc9dec
Output folder: ./output/
Tool for initial candidate selection: blast
Tool for tree construction: fasttree
CPUs for BLASTp: 4
CPUs for RAxML: 4
Reference MYB file: /xxx/myb/AthRefMYBs.txt	abc8047bebc188b8a20935fe82467b5d
Type of input: PEP
blastp path: blastp
makeblastdb path: makeblastdb
mafft path: mafft
raxml path: raxml-ng
fasttree path: fasttree
Minimal BLASTp hit similarity cutoff: 60.0
Maximal number of BLASTp hits per bait: 100
Minimal BLASTp hit alignment length: 75
Number of neighbours to consider in classification: 15
Factor for branch length cutoff in identification of neighbours: 10.0
Minimal number of neighbours required for classification as MYB: 0
Distance cutoff for paralog clade masking: 10.0
MAFFT version: v7.505 (2022/Apr/10)
FastTree version:  2.1.11 
RAxML version: RAxML-NG v. 0.9.0 released on 20.05.2019 by The Exelixis Lab....
hmmsearch version detection failed.
