## Additional file 4 for "Genome sequence and RNA-seq analysis reveal genetic basis of flower coloration in the giant water lily *Victoria cruziana*"

bHLH_annotator.py version: v1.04
bHLH bait file: ../../bHLH_annotator/data/bHLH_baits.fasta	cf967906cd921ac579e1514fc81c6eca
bHLH bait info file: ../../bHLH_annotator/data/bHLH_baits.txt	454e90dfcdcb2e82ea663639a96f47d6
 thinned bait file: ../../bHLH_annotator/data/bHLH_baits_optimised.fasta	0c9e29a40a8e299c5300f77f6c43d614
Subject FASTA file: ./proteins.fasta	73d58054958954b1d221fcf612bc9dec
Output folder: ./
Tool for initial candidate selection: blast
Tool for alignment: mafft
Tool for tree construction: fasttree
Maximal CPUs for ingroup/outgroup classification: 1
CPUs for BLASTp: 1
CPUs for RAxML: 1
HMM bait motif file: ../../bHLH_annotator/data/bHLH_baits_optimised.hmm	b7ffe237696c182a22e0fe725fe1701f
Reference bHLH file: ../../bHLH_annotator/data/AthRefbHLHs.txt	3dea98bdc522a4e0df4df07b2f9d5d34
Ath family members file: ../../bHLH_annotator/data/bHLH_at.fasta	c001f05e26bf4d4e3ae1424eee0dcd74
Motif search file: ../../bHLH_annotator/data/bHLH_motifs.hmm	c38f1a05621c6a40bd5cfed9d133f66e
Type of input: PEP
Prefix of output file names: 
Preventing of splitting sequence names at first space (--keepnames): True
Reducing paralogs to one representative (--collapse): False
blastp path: blastp
makeblastdb path: makeblastdb
hmmsearch path: hmmsearch
mafft tool path: mafft
muscle tool path: muscle
raxml path: raxml-ng
fasttree path: fasttree
Minimal bitscore cutoff: 60
Minimal BLASTp hit similarity cutoff: 40.0
Maximal number of BLASTp hits per bait: 100
Minimal BLASTp hit alignment length: 80
Minimal score for candidate to be considered as ingroup: 0.5
Number of neighbours to consider in classification: 10
Factor for branch length cutoff in identification of neighbours: 5.0
Minimal number of neighbours required for classification as bHLH: 0
Filter candidates by hmm motif: False
Distance cutoff for paralog clade masking: 10.0
Parallelization at ingroup/outgroup classification: False
Maximal number of candidates per parallel ingroup/outgroup classification: 200
Muscle version: 
MAFFT version: v7.505 (2022/Apr/10)
FastTree version: PLEASE_ADD_MANUALLY
RAxML version: ...
hmmsearch version:  HMMER 3.4 (Aug 2023); http://hmmer.org/\n...

Execution time: 0h:21min:33sec
