## Additional file 5 for "Genome sequence and RNA-seq analysis reveal genetic basis of flower coloration in the giant water lily *Victoria cruziana*"

**Principle component analysis of young and mature samples**

Principal Component Analysis of the RNA-seq samples confirming the clustering of biological replicates in their respective groups. Young petals are represented in A, mature petals in B.

**A**

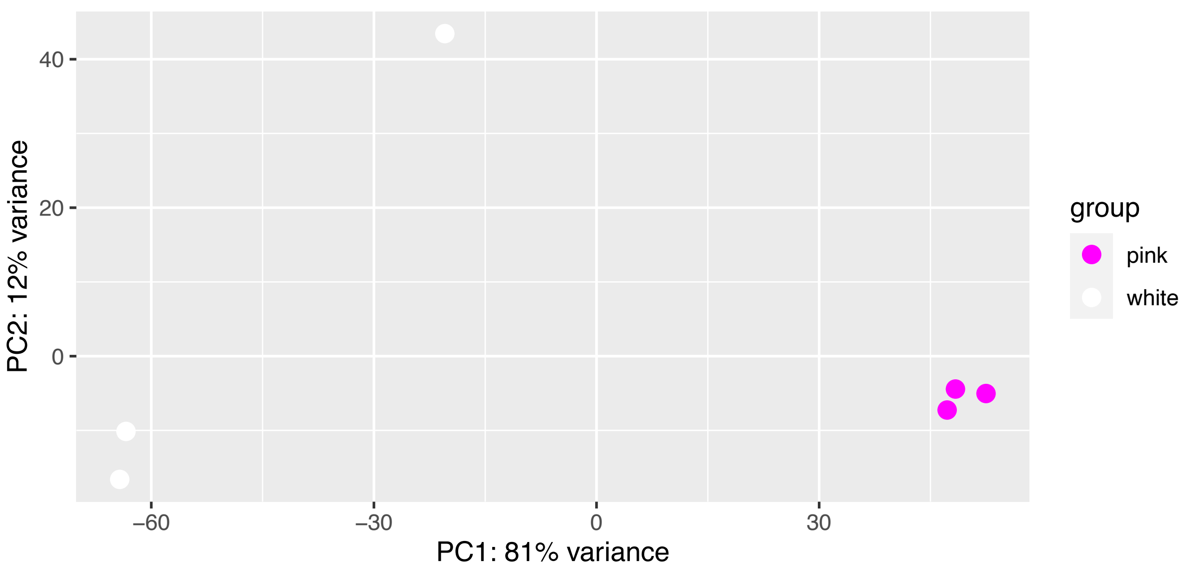


**B**

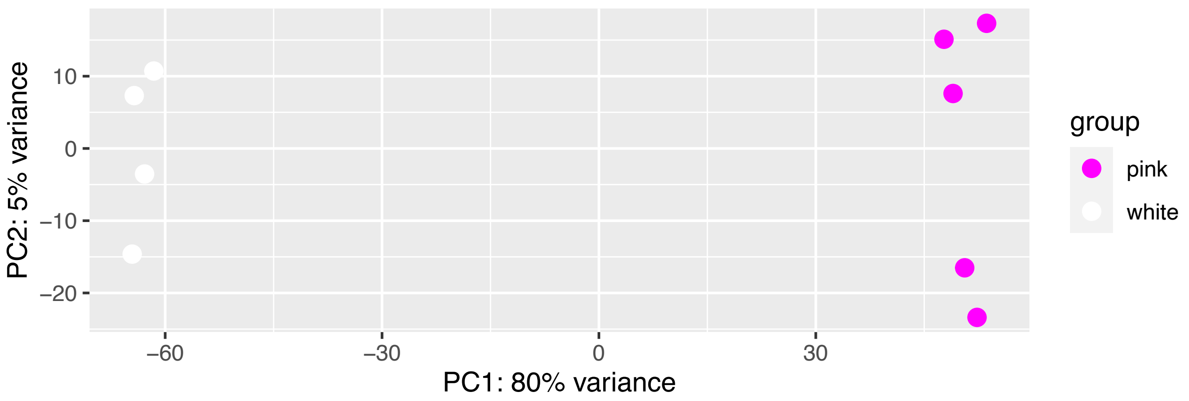
