## Supplementary figures and images for "Genome sequence and RNA-seq analysis reveal genetic basis of flower coloration in the giant water lily *Victoria cruziana*"

### Additional file 8

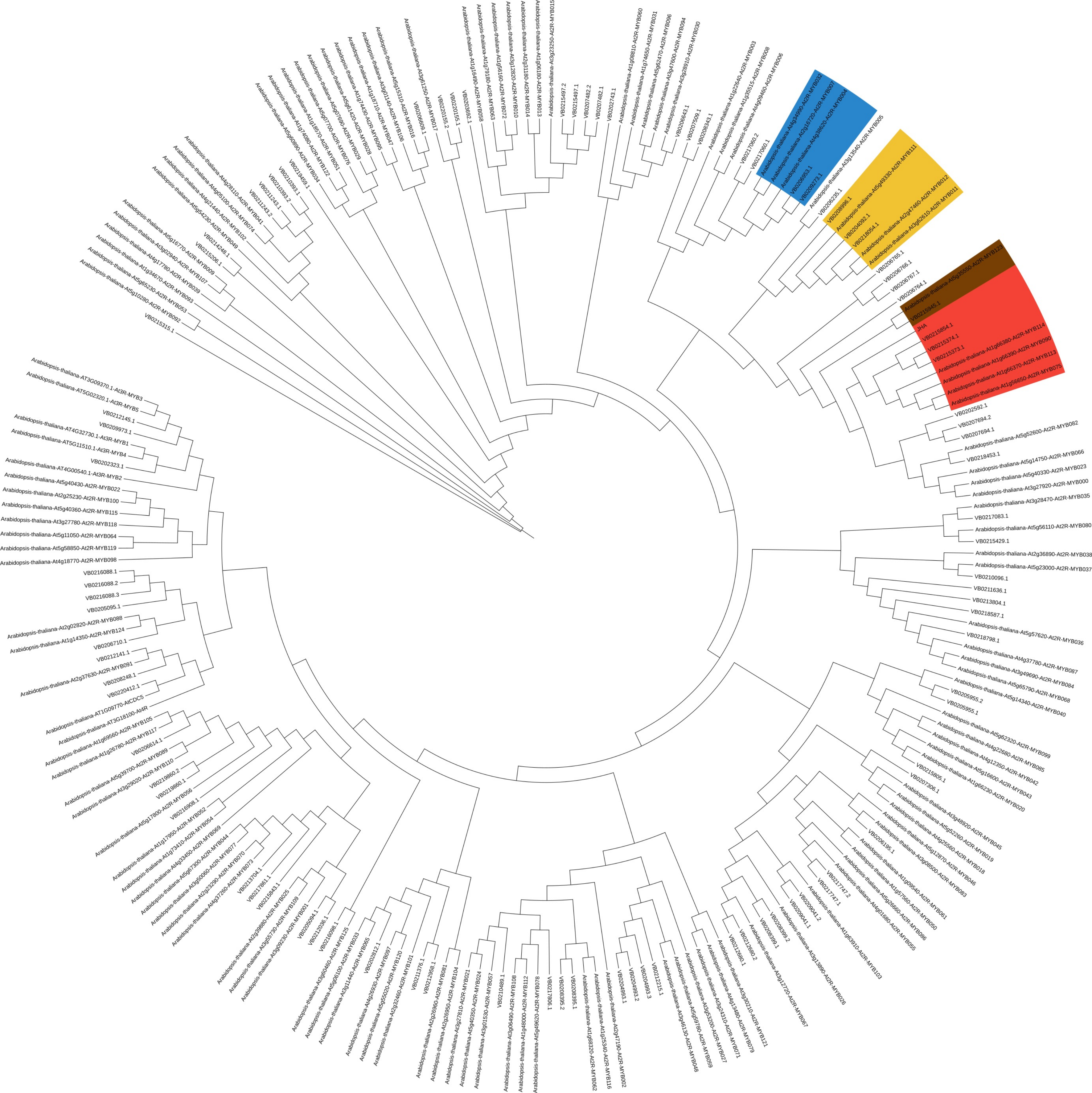
