## Additional file 10 for "Genome sequence and RNA-seq analysis reveal genetic basis of flower coloration in the giant water lily *Victoria cruziana*"

| Replicate | Flower closed | Flower open: front | Flower open: back |
|---|---|---|---|
| 1 | 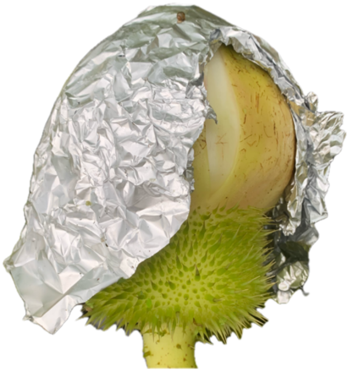 | 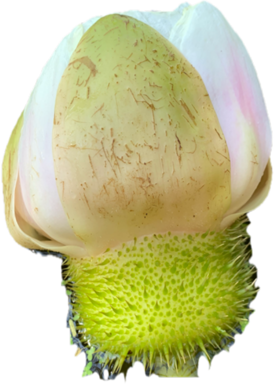 | |
| 2 | 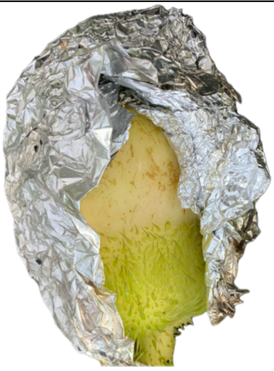 | 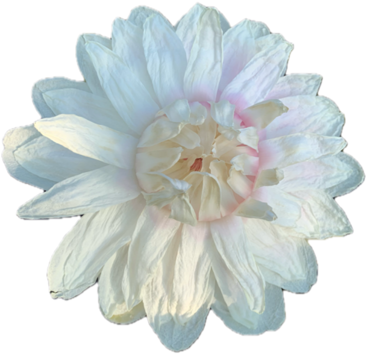 | 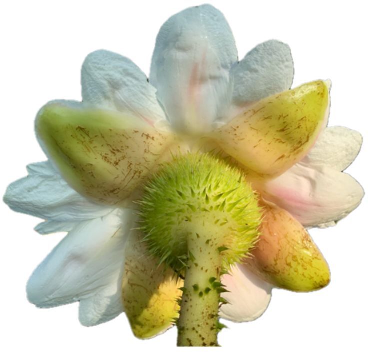 |
| 3 | 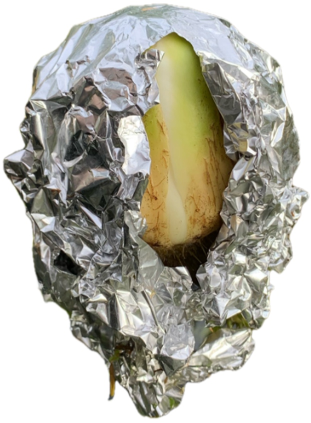 | 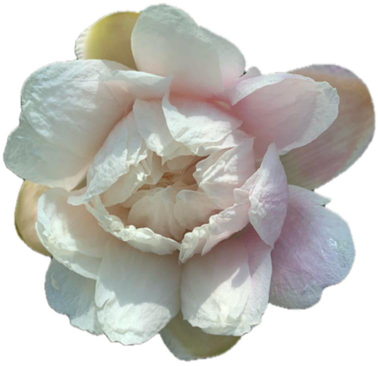 | 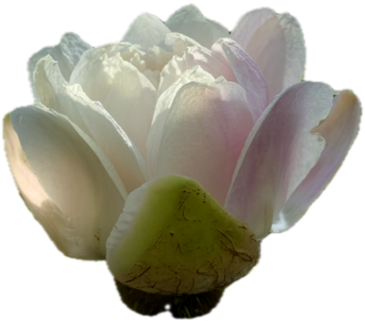 |
| 4 | 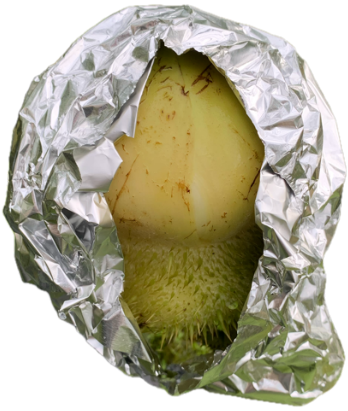 | 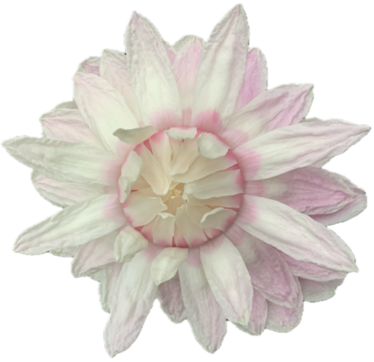 | 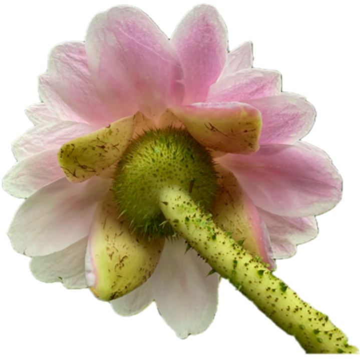 |

**Table:** Visualization of flowers after sunlight exposure experiment. Flower buds were wrapped into aluminum foil under water as soon as they emerged. After perception of fruity smell, a small part of the aluminum foil has been cut out. This enabled sunlight exposure to a defined area of the flower bud. After approximately one day, flowers were examined regarding coloration pattern. All stages were documented by images as presented above. Sunlight exposure experiments have been conducted during July – September 2024.
